# Nonequilibrium bending fluctuations reveal microtubule mechanics *in-vivo* and their regulation by glutamylation

**DOI:** 10.1101/2023.10.26.564252

**Authors:** Kengo Nishi, Sufi Raja, An Pham, Ronit Freeman, Antonina Roll-Mecak, Fred C. MacKintosh, Christoph F. Schmidt

## Abstract

Cells can be described as active composite materials. The mechanical properties of cells are controlled by complex polymer networks and are dynamically tuned for diverse cellular processes driven by force-generating motor proteins. Microtubules are the most rigid protein polymers in the cytoskeleton, and their material properties have been measured *in vitro* by active bending or by analyzing thermal bending fluctuations. Microtubule mechanics in living cells are extremely difficult to probe directly, while fluctuations are difficult to interpret because they are generated by active forces in a surrounding cytoplasm with poorly understood material properties. Here we introduce a method to measure the elastic properties of microtubules in living cells by making use of motor-generated forces that drive bending fluctuations. Bending dynamics are governed by three main factors: microtubule material properties, cytoskeletal active forces, and the response characteristics of the surrounding cytoplasm. We show theoretically that, when one factor can be independently determined, the other two can be derived from observed fluctuations. Using this method we discovered that polyglutamylation, a post-translational modification enriched on microtubule arrays that need to withstand large mechanical forces such as those in axons or cilia, increases microtubule stiffness in living cells. Our work provides a theoretical and experimental framework to study microtubule mechanics and their regulation by tubulin modifications and microtubule effectors in complex cellular environments. The approach can be extended to other large aspect ratio cellular structures such as the endoplasmic reticulum or the mitochondrial network.

## Introduction

Evolution has produced highly complex and versatile functional biomaterials. These materials are typically active, since all life processes need constant energy dissipation. Based on a universal set of molecular components including lipids, proteins, nucleic acids, and polysaccharides, cells generate dynamic and adaptable functional structures by polymerizing and depolymerizing, connecting, modifying, and adapting structural building blocks, many of which are acted on by force-generating mechanoenzymes. Microtubules (MTs) are one of the main structural components of the cytoskeleton. MTs are tubular, linear polymers constructed by the dynamic non-covalent association of *αβ*-tubulin heterodimers, and they support diverse cellular processes, including meiosis and mitosis, transport and signaling, as well as assembly of cilia and flagella for movement [1]. To optimally serve these diverse functions, chemical and mechanical MT properties are fine-tuned by post-translational modifications (PTMs) of *αβ*-tubulins [2-5]. Glutamylation, the addition of glutamates to the C-terminal tails of tubulin, is evolutionarily widespread from ciliates to humans, and is one of the most abundant modifications in the human brain [6]. This modification is introduced by a large family of glutamylation enzymes, the tubulin tyrosine ligase like family (TTLL), with important functions in neuronal development and axonemal assembly [6]. In cells, glutamylation is found on microtubules with slow turnover such as those in axons, cilia and parts of the mitotic spindle [7-11]. The consequences of this PTM for microtubule mechanics, however, remain unknown. The elastic properties of microtubules have been mostly studied in *in vitro* experiments, using reconstituted microtubules, either by observing thermal bending fluctuations and applying the fluctuation-dissipation theorem (FDT) [12-14] or by active manipulation [15-24]. There remains order-of-magnitude disagreement between results, presumably due to structural defects introduced during reconstitution of microtubules and/or interactions with other proteins [12, 25-30]. Furthermore, it is still unclear what the precise mechanical properties of microtubules are in the living cell. *In vivo* experiments are complicated by the structural complexity of cells. In addition, living cells are not in thermodynamic equilibrium. Force generators, such as myosin motors, hydrolyze adenosine triphosphate (ATP) and cause non-equilibrium fluctuations. Such active forces drive bending and buckling dynamics of microtubules [31-34]. Observed fluctuations alone are thus not sufficient to determine the mechanical response properties of microtubules as long as both the active forces from the surrounding cytoplasm and the mechanical response properties of the cytoplasm are unknown [33, 35, 36]. Several studies have applied external forces to living cells to measure global and local response using atomic force microscopy (AFM), optical tweezers (OT), and magnetic tweezers (MGT) [36-46]. Such experiments, however, are not sensitive to the mechanical properties of microtubules.

Here, we propose a new approach to determine the material properties of individual microtubules in living cells by analyzing their bending fluctuations. Non-muscle myosin II aggregates are the dominant force generators driving random fluctuations as well as collective motions of the actin cytoskeleton in most non-muscle cells [33, 36]. These forces also drive fluctuations of the embedded microtubules (Fig. 1A) [33]. Microtubule bending dynamics are governed by the balance of four forces: the elastic restoring force due to bending, the viscoelastic resistance from the surrounding cytoplasm, the thermal forces, and the active forces. We have developed a theoretical model to relate microtubule bending fluctuations to those four forces. Using the model, we can estimate the active forces and the bending stiffness of microtubules once cytoplasmic response is measured independently. The latter was achieved by applying force steps to magnetic probe particles embedded in the cell cytoplasm (Fig. 1B). We also used prior knowledge about the spectral characteristic of myosin-actin generated forces in the cell [33, 36, 47]. To validate the approach, we used the fact that the cytoplasmic elastic response can also be estimated from the shape of localized large-amplitude bends of microtubules under certain conditions (Fig. 1C). We found that the measured *in vivo* microtubule stiffness falls in the range of published *in vitro* data. Furthermore, we show that overexpression of the glutamylase TTLL6 increases microtubule bending stiffness in cells.

**Figure 1.**
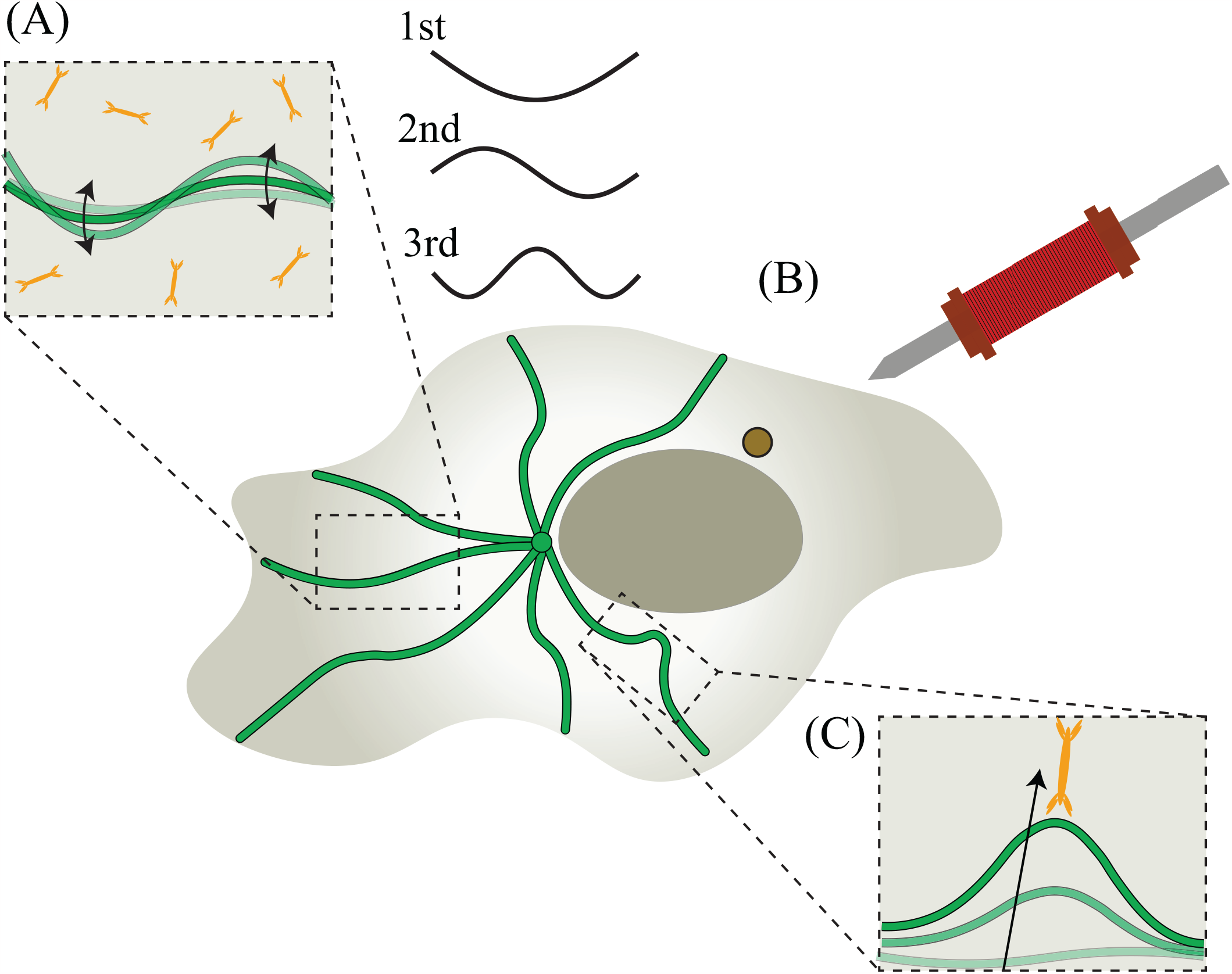
Schematic illustration of microtubule bending dynamics in cells. **(A)** Molecular motors and other active processes provide a (typically) incoherent background of fluctuating forces, driving transient bending deformations of microtubules that we record by confocal fluorescence microscopy. The shapes of the fluctuating microtubules are decomposed into Fourier modes and the first three modes are visually presented. **(B)** Schematic of magnetic tweezers microrheology to measure cytoplasmic viscoelasticity in HeLa cells. The gradient force generated by an electromagnet with a ferromagnetic core ending in a sharp tip near the cell acts on 1 *μ*m super-paramagnetic beads electroporated into the HeLa cells. **(C)** Occasionally, molecular motors produce strong isolated forces causing sharp localized microtubule bends.

## Results and Discussion

We used HeLa cells as a model system and imaged fluorescently labeled microtubules using SiR-tubulin [48] in a confocal microscope (Fig. 2A), described in detail in the Supporting Information (SI). Although microtubules have a persistence length of ∼ 1 mm at room temperature and therefore show only weak thermal bending fluctuations on the scale of micrometers [12, 25, 49], microtubules exhibit significant curvature in living cells. Such strong bending is mainly caused by nonthermal forces such as acto-myosin contractility or the dissipative polymerization of microtubules [31-34]. Bending on length scales ≳ 6 μm can also result from slow, large-scale cellular restructuring, such as the retrograde flow of the entire cytoskeletal network [31]. Such long-wavelength bends relax extremely slowly because they are mechanically constrained by the surrounding viscoelastic cytoskeleton. Therefore if one wants to observe full relaxation and sample uncorrelated fluctuations on practical recording time scales (minutes), one needs to focus on length scales shorter than ∼ 6 μ(Fig. 2B). To evaluate microtubule mechanical properties from these non-equilibrium bending fluctuations, we quantified and analyzed scale-dependent power spectral densities (PSDs) of microtubule deflection amplitudes. We tracked the shapes of fluorescently labeled microtubules from confocal fluorescence microscopy recordings of substrate-adherent cells (Fig. 2A). A time series of snapshots of conformations of an individual microtubule is shown in Fig. 2B. The instantaneous contours were quantified by the transverse deflection *u*(*s*,*t*) from a line connecting the ends (not including center-of-mass movement and global rotation) and then decomposed into Fourier modes as a sum of sines:

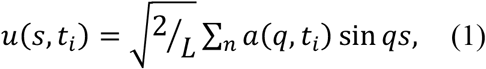

with the contour length variable *s* (see SI) and the discrete times *t*_*i*_ at which frames were recorded. The bending mode amplitudes *a*(*q*,*t*_*i*_) parametrize the shape of filaments for a set of discrete wave vectors *q*_*n*_ =*nπ*/*L* =π/*L*_eff_, where *n* = 1,2,3… is the mode number. *L* and *L*_eff_ are the fiber length and effective mode length, respectively. The characteristic axial extent of a bend caused by a lateral or axial point force is governed by the competition between the bending energy of the filament and the dynamic shear elastic modulus of the cytoplasm. Thus, we can quantify the modulus of cytoplasm from the characteristic length of localized bends once the microtubule stiffness is known or *vice versa*.

**Figure 2.**
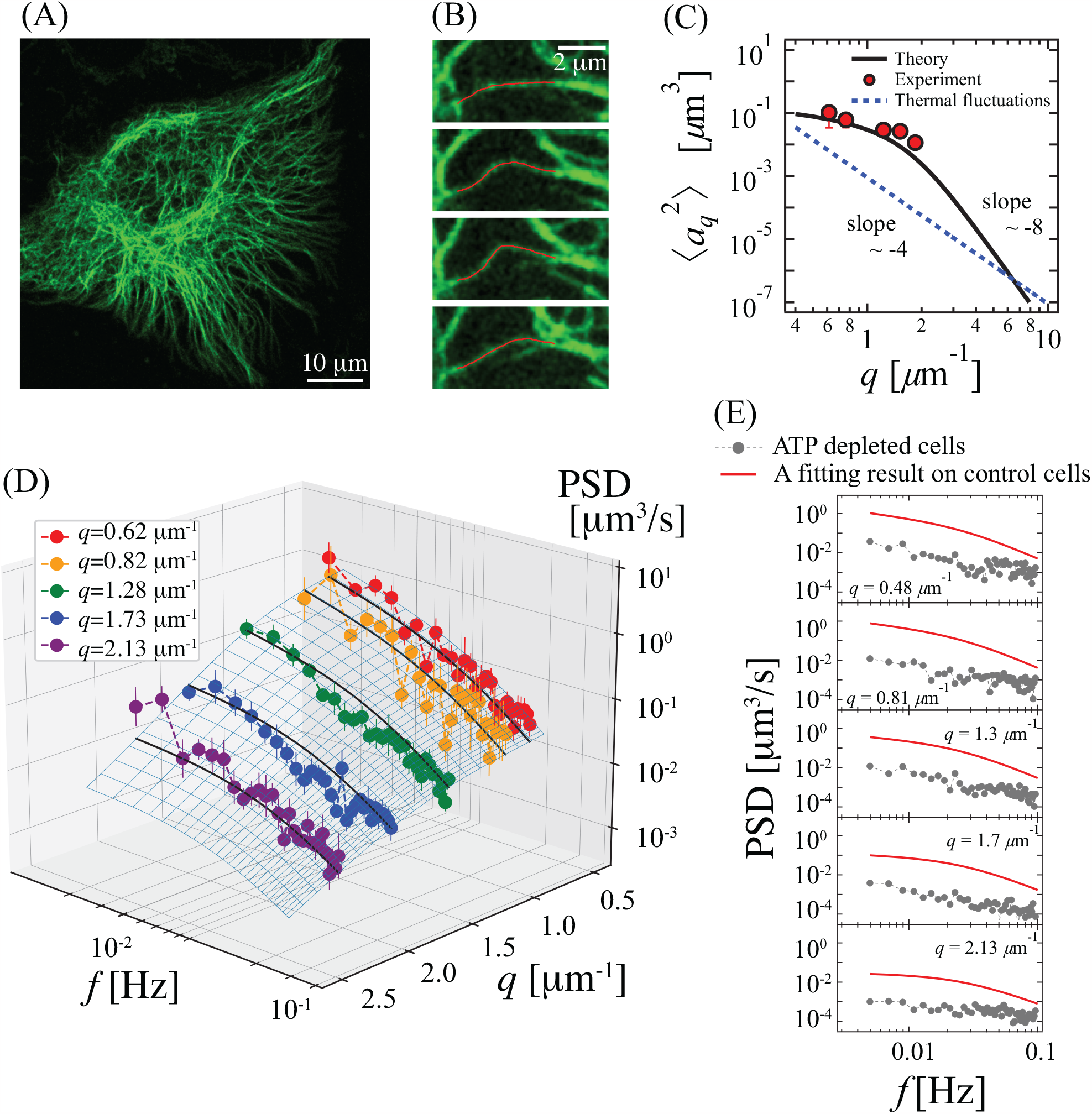
Bending fluctuations of microtubules in living cells. **(A)** A substrate-adherent living HeLa cell with microtubules fluorescently labeled using SiR-tubulin (green) (see Supplementary movie S1). **(B)** Microtubules undergo significant bending fluctuations caused by nonthermal forces. Consecutive images are separated by 20 sec. The red overlay is the filament contour approximated by image analysis. **(C)** Variance of the mode amplitudes, plotted vs. wave number of 18 microtubules. The mode amplitude variance (red circles) (average of 18 microtubules) is significantly larger than the predicted thermal fluctuations: ⟨Δa_0_^2^⟩ = 1/(*l*_p_*q*^4^) with a persistence length of *l*_p_ ∼ 1 mm (blue dashed line), using a value reported value from *in vitro* studies [25]. Measured variances are consistent with the theoretical expectation (black line) applying the Wiener-Khinchin theorem and using fitting results with Eq. (6) to PSDs shown in (D). **(D)** Global fit of mode-amplitude power spectral densities of microtubules in Hela cells. Binned PSDs from 18 microtubules at *q* = 0.62, 0.82, 1.28, 1.73, and 2.13 *μ*m^-1^ as plotted as circles. Error bars are ± SEM. The bin widths of each *q* point are approximately ∼ 0.3 *μ*m^-1^. A 2D fitting result of PSDs with Eq. (6) is shown (gray mesh plane). Black solid lines represent slices of the fitted plane at *q* = 0.62, 0.82, 1.28, 1.73, and 2.13 *μ*m^-1^, respectively. **(E)** For comparison, we tracked and analyzed bending fluctuations of four microtubules in ATP depleted cells. Their mode amplitude fluctuation PSDs (gray circles) were smoothed by binning in *q*-space at *q* = 0.48, 0.81, 1.3, 1.7, and 2.1 *μ*m^-1^. Also plotted are 1D slices of the 2D fit surface to the mode amplitude fluctuation PSDs of microtubules in control cells at *q* = 0.48, 0.81, 1.3, 1.7, and 2.1 *μ*m^-1^ (red line). Four microtubules from two different ATP-depleted cells were tracked.

To obtain microtubule bending stiffness in living cells, we first calculated amplitude variance as a function of mode number ⟨a^2^(q)⟩ (Fig. 2C) from recordings either in the cell periphery or beneath the nucleus of surface-adherent cells. Fourier amplitudes were calculated frame-by-frame from recordings taken at frame rates of 0.25 ∼ 0.5 Hz. To quantify the average dynamics, we calculated the mean-squared amplitude difference as a function of lag time, 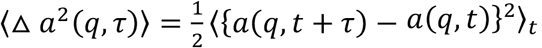 . At long lag times *π*, microtubule shapes will become uncorrelated from their initial shapes, leading to ⟨Δ a^2^(q, τ → ∞)⟩ = ⟨a^2^(q)⟩.

For the case of thermal equilibrium, the equipartition theorem predicts that ⟨a^2^(q)⟩ = *k*_B_*T*/(κ*q*^4^) = 1/(*l*_p_*q*^4^) with the thermal persistence length *l*_p_ = κ/*k*_B_*T* that is on the order of millimeters [12, 25] (Fig. 2C). We found that the bending amplitude variance of microtubules in cells was significantly larger than the predicted thermal fluctuations, consistent with previous reports [31]. To further analyze these bending fluctuations, we calculated power spectral densities (PSDs) ⟨|a(q, ω)|^2^⟩ of individual mode amplitudes (Fig 2D). Consistent with the mode-number dependence of the variances (Fig. 2C), we find that the PSDs decrease in amplitude with increasing *q* due to the bending elasticity of the microtubules. Note, however, that the functional form of the PSDs also changes with *q*. In addition, we observed a consistent and significant drop in PSD amplitudes for all modes in ATP depleted cells (Fig. 2E). This large drop confirms that active forces dominate the bending fluctuations of microtubules for the observed frequencies < 0.1 Hz, corresponding to times longer than a few seconds. To extract microtubule material properties from the measured PSDs, we model the dynamic bending fluctuations of a slender filament driven by active forces and embedded in a viscoelastic continuum, the cytoskeleton. We assume that force-generating motors are randomly distributed through the medium and that both thermal forces and motor forces propagate through the medium and induce transverse fluctuations of microtubules. We describe transverse filament motion by a Langevin equation for the force per unit length on a filament segment at position *s* [50]:

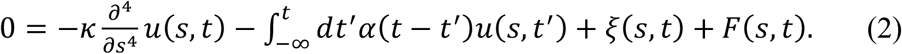

The first term accounts for the elastic restoring force [12] due to the filament bending rigidity *κ*. The second term is the viscoelastic drag from the embedding medium, where the resistance per unit length is given by the memory function *α*(*t*). For the time and length scales relevant to this work, an inertial term on the left-hand side of Eq. (2) should be negligible and has been set to zero. The Fourier transform of the memory function *α*(ω) is proportional to the complex shear modulus of the medium, *G*(ω) with *α*(ω) ≈ 4*πG*(ω)/ln(*AL*_eff_/*d*) as shown previously [50], where *A* ≅ 2.3 and *d* is the diameter of the filament. The stochastic term ξ(*s*,*t*) models the net thermal force exerted on the filament. In order to account for the impact of active forces produced by motors located in the vicinity of the filament, we add the active force term *F*(*s*,*t*).

We analyze the bending dynamics of the filaments by expanding *u*(*s*,*t*) into spatial Fourier modes as approximations of orthogonal dynamic eigenmodes (Eq. 1). Using the decomposition in Eq. (1), we can write Eq. (2) as:

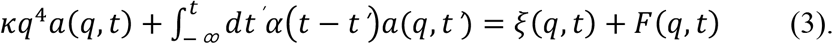

The temporal Fourier transform of this allows us to relate the amplitudes *a* to the forces via *a*(*q*,*ω*)=*χ*(*q*,*ω*)[ξ(*q*,*ω*)+*F*(*q*,*ω*)] and the response function χ(*q*,*ω*)=[κ*q*^4^+*α*(*ω*)]^-1^. Inside the cytosol, the thermal force ξ(*s*,*t*) can be neglected because active forces *F*(*q*,*t*) are expected to be dominant over ξ(*q*,*t*) for fluctuations slower than ∼10 Hz, the regime we focus on in this study, as shown in previous studies [36, 37]. With these assumptions, we obtain the power spectral density (PSD) of mode amplitudes as:

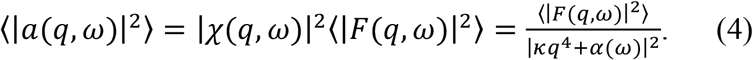

To interpret measured PSDs, we first need to approximate the scale-dependent force spectrum, ⟨|*F*(q, ω)|^2^⟩ which is the Fourier transform of the autocorrelation of active forces, ⟨*F*(*s, t*)*F*(*s*′, *t*′)⟩. We consider motors that are located near the filament since those are expected to most strongly drive the filament fluctuations. We assume that force generation by different motors is uncorrelated. As widely observed in the cytoskeleton and in reconstituted motor-activated cytoskeletal model systems [35, 36, 47], the temporal force correlation function at a point in the system resulting from motor activity decays exponentially. Taken together, this will cause the correlation of active forces to have a functional form of: 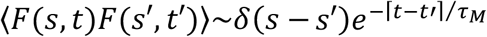. Thus, the Fourier transform of the correlation of active forces is a Lorentzian:

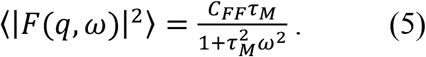

Here, τ_M_ is the correlation time of the force generators, which has been reported to be ∼10 s for actomyosin activity in the cytoskeleton [36]. *C*_*FF*_ is a constant and has units of a line density of squared-force [N^2^/m]. Based on previous studies [51, 52], *C*_*FF*_ is approximated as *C*_*FF*_∼ ⟨*f*_0_^2^⟩,/*l*_*M*_ in the limit of high densities of molecular motors, where ⟨*f*_0_^2^⟩ and *l*_M_ are the mean squared active forces on a single point and the distance between neighboring motors, respectively. Combining eqs. (4) and (5), we find for the PSD of the actively driven, fluctuating filament:

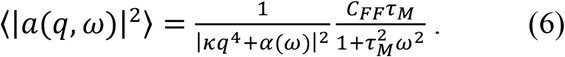

This result relates the PSDs of the eigenmode amplitudes to the relevant physical properties of the system, i.e. the active force fluctuations *C*_*FF*_ and τ_M_, the bending stiffness of the filaments, and the viscoelasticity of the embedding medium *α*(*ω*). As *q* increases, the effect of bending stiffness becomes prominent in the PSDs, and therefore high-*q* PSDs need to be included in the data evaluation in order to estimate the filament bending stiffness. In principle, one could attempt to obtain all five unknown parameters in Eq. (6), including the function *α*(*ω*) by a global fit of all measured PSDs. Resulting uncertainties would be large, however. To measure the bending rigidity of microtubules in living cells more accurately, we thus first independently determined the viscoelasticity of the cytosol, *G*(*ω*), by performing active microrheology [35-37, 44, 53]. Previous studies of active microrheology applied oscillatory forces using optical traps (OT) [36, 37, 53]. However, in an OT experiment, intense laser light can induce photodamage in cells, and measurements are easily perturbed by refractile objects other than the probe particles in the cells, such as the nucleus and various organelles. To avoid these potential problems here, we used “magnetic tweezers”. We embedded 1 μm-diameter magnetic beads in the cytosol of Hela cells (Fig. 3A) and applied magnetic force steps on these probe particles (Fig. 3B). From the recorded displacement response, *d*(*t*), of the beads we calculated creep compliance *J*(*t*) = σ(*t*)/*α*_0_ = 6*ad*(*t*)/*f*, where *a* is the bead radius, σ(*t*) is the time-dependent shear strain, *α*_0_ the applied constant stress, and *f* is the applied magnetic force. We found that the creep compliance shows a power-law behavior, *J*(*t*) ∝ *t*^β^ with *β* = 0.45 (± 0.01) (Fig. 3C), indicating a broad spectrum of relaxation times in the cytoplasm as commonly observed in previous studies [37]. Viscoelastic response can also be expressed in terms of the frequency-dependent shear modulus *G*(*ω*), related to the time-dependent compliance through a Fourier transform:

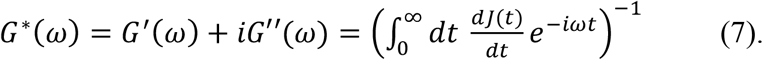

This equation is the Fourier transform of 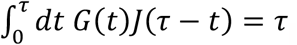, which relates the relaxation modulus *G*(*t*) = *α*_0_/γ(*t*) to the creep compliance by a convolution [54, 55]. We applied the five-point stencil method [56] to accurately calculate the numerical derivative, and Simpson’s rule for the subsequent integral [57]. We find that the dynamic shear modulus varies with frequency as a weak power-law with an exponent in the range of 0.3 - 0.5. This slow stress relaxation has been widely observed in living cells [36, 37, 40, 44-46, 58] and may be related to soft glassy dynamics or transient cross-linking by cytoskeletal networks. We find good agreement between *G*_M_(*ω*) that we measured with magnetic tweezers with *G*_OT_(*ω*) reported in the literature [37] (Fig. 3D).

**Figure 3.**
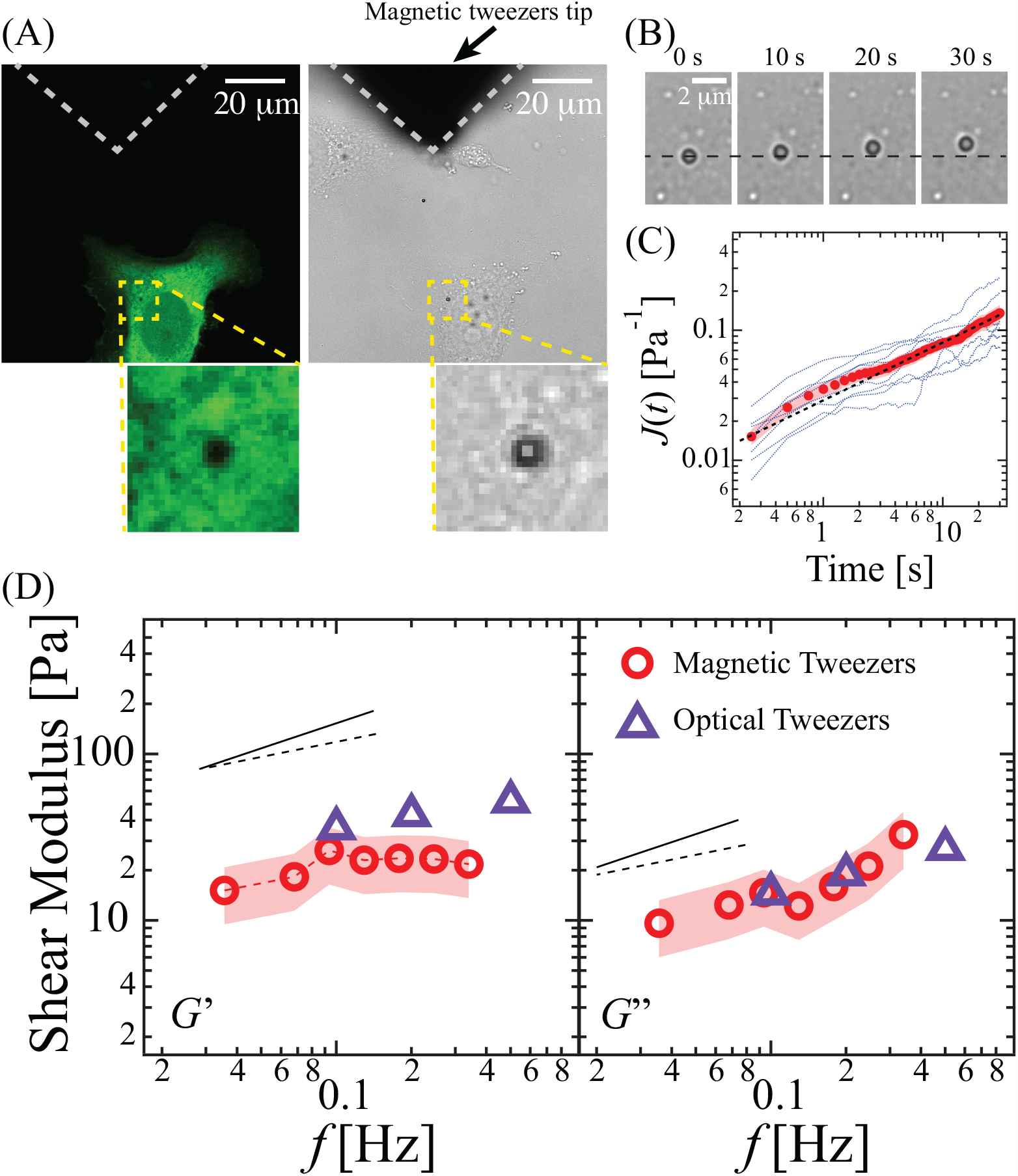
Probing cytoplasmic viscoelasticity by magnetic tweezers. **(A)** Confocal and brightfield images of magnetic tweezers experiment in a substrate-adherent HeLa cell expressing GFP. The dark spot in the fluorescent image at the bead position proves that the bead is successfully introduced in the cytoplasm. The tip of the electromagnet is 45 ∼ 50 *μ*m away from the magnetic bead and applies ∼ 60 pN force for 30 s (see Supplementary movie S2). **(B)** Time sequence of brightfield images of typical bead displacements towards the tip of the magnet in the cytoplasm with ∼ 60 pN force. Beads move ∼ 500 nm in 30 sec, a small enough distance to avoid nonlinear response or rupture. **(C)** Displacement is converted to creep responses (*J*(*t*)) (9 measurements from 9 different cells). Individual response curves (blue), average (red) and powerlaw fit (dotted black line). The applied force *f* is assumed to be constant since the displacement of the beads (∼ 500 nm) is negligible compared to the distance between the tip and the bead. **(D)** Complex shear modulus of the Hela cell cytoplasm measured from the creep response to force steps (open circles). Creep functions are converted to the complex shear modulus using Eq. (7), real part or storage modulus *G’* and imaginary part or loss modulus *G”*. Errors are ± SD. For comparison, values obtained from oscillatory experiments using optical tweezers were taken from Ref. [37]. Slopes of 0.3 and 0.5 are also plotted to guide the eye.

To now obtain the bending stiffness of microtubules in the living cells, we globally fitted the scale-dependent PSDs (Fig. 2D) with a single 2D plane defined by Eq. (6) with just three free parameters. *C*_*FF*_, τ_M_, and *κ*. Using the magnetic-tweezers rheology results, we fixed the memory functionα(*ω*) = *k*_0_*G*(*ω*) using 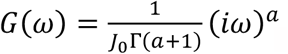, where *J*_0_ and *a* are obtained from a power-law fit to the rheology data, *J*(*t*) = *J*_0_*t*^a^,. and *Γ* is the gamma function. From the global fit to five scale-dependent PSDs evaluated from 18 microtubules, we find κ= 4.1 (± 0.7) ×10^-24^ Nm^2^,τ_M_ = 5.9 (± 0.7) s, and *C*_FF_ = 4.8 (± 1.3) ×10^1^ pN^2^/μm. Uncertainties in the fitting parameters come from the variations in the measured PSDs from different microtubules and in the complex shear modulus measured by active magnetic-bead microrheology and were estimated by a bootstrapping method (see SI). Assuming *C*_FF_ ≈ ⟨*f*_0_^2^⟩/*l*_M_, where *f*_0_ is an active force of ∼ 7 pN, the line density of motors *l*_M_ is estimated to be on the order of ∼ 1 μm^-1^, which is consistent with imaging data of non-muscle myosin bundle densities in cells [59]. The parameter τ_M_ found in this study is similar to the value reported in the literature [36]. The persistence length, *l*_p_ =κ/*k*_B_*T*, of the microtubules we tracked in cells comes out to be 1.0 (± 0.2) mm, which lies in the range of values measured in aqueous buffers and actin solutions [12, 18, 25]. This consistency validates our new method of determining the material properties of microtubules in living cells from active shape fluctuations.

To check for consistency, we evaluated the total mode variance from the scale-dependent PSDs (Fig. 2C). We used the 2D fitting plane to the PSDs (Fig. 2D) to obtain fitting parameters(*κ* = 4.1 ×10^-24^ Nm^2^,τ_M_ = 5.9 s, and *C*_FF_ = 4.8 ×10^1^ pN^2^/μm). Subsequently, we applied the Wiener-Khinchin theorem, 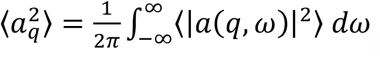, to calculate the total variance for each mode (black solid line in Fig. 2C). We find that experimental data (red circles) fits with our theoretical prediction within the observed *q*-range, demonstrating internal consistency of our analysis.

Up to here, we have calculated microtubule stiffness after using an independent method to determine cytoplasmic response. Reversing this logic, we can also use the dynamics of microtubules, once we know their bending stiffness in cells, to probe the cytosol mechanics. This approach makes it possible to monitor intracellular mechanical response characteristics without inserting foreign particles as mechanical probes, as long as the forces are well-defined. In the approach we introduce in the following, we fit the shapes of strongly bent microtubules. In living cells, some microtubules show deformations that don’t relax on a time scale of minutes [31], reflecting strong constraints in the cytoskeleton. We avoided such microtubules by selecting localized bends on microtubules which were initially straight and then sprang back to a straight shape after a transient deformation (Fig. 4A). Note that in this approach, the magnitude of the force does not need to be known, but we need to be sure that it is a point force acting laterally on a given microtubule. It turns out that microtubules in cells occasionally experience localized large-amplitude transverse bends that are likely caused by nearby myosin motor complexes exerting an approximately point-like force. Similar deformations have been observed in response to compressive forces from the cell boundary exerted on growing microtubules, leading to Euler buckling [25, 32]. Fitting the shapes of such strongly bent microtubules can be used to quantify the zero-frequency elasticity of the embedding cytoskeleton. The axial extent of a bend is set by the minimization of total elastic energy, balancing the bending energy of the filament against the elastic energy of the deformed embedding cytoskeleton [25]:

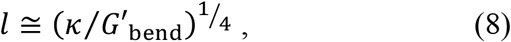

with the bending rigidity of the microtubule, *κ*, and the zero-frequency shear elastic modulus of the cytoskeleton, *G*’_bend_. *G*’_bend_ can thus be calculated from the observed length scale *l*. The bend length *l* can be extracted from microscopic images by fitting bent shapes using the following equation [25],

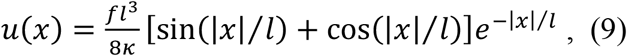

where *f* is the applied point-like force in the middle of the bend, *u*(*x*) is the amplitude of the localized bend as a function of position *x* along the filament. To make sure that the bends we observed were not caused by Euler buckling, we selected localized bends at least 4 *μ*m away from the tip of microtubules since Euler buckling is caused by axial compression at the ends of growing microtubules. Fig. 4B shows that the shapes of strongly bent microtubules we observed are well described by Eq. (9), which confirms the assumption that the dominant force acting was a point force. We tracked 11 microtubules, fitted them with Eq. (9) (Fig. S2), and extracted a bend length *l* = 0.64 (± 0.1) *μ*m (Fig. 4C). With the value of *κ* = 4.1 ×10^-24^ Nm^2^ obtained above, we find for the cytoplasm elasticity *G*’_bend_ = 24 (± 17) Pa. The good agreement between the real part of *G*_M_(ω) and *G*’_bend_ demonstrates the consistency of the methods, and shows that the cytoskeletal elastic response can be successfully estimated from the shape of localized transverse microtubule bends. Cytoskeletal elasticity in the periphery of the surface-adherent HeLa cells presumably largely stems from actin networks. To confirm this, we pretreated Hela cells with 1.5 *μ*M latrunculin A for 15 min to (partially) disrupt the actin network. We found that the wavelength of localized microtubule bends increased to 1.06 (± 0.15) *μ*m in latrunculin A-treated cells, from 0.64 (± 0.1) *μ*m in untreated cells (Fig. 4C). The elastic modulus obtained from those localized bends in latrunculin A-treated cells again roughly agrees with the real part of *G*_M_(ω) measured by magnetic tweezers in cells treated in the same way (Fig. 4D).

**Figure 4.**
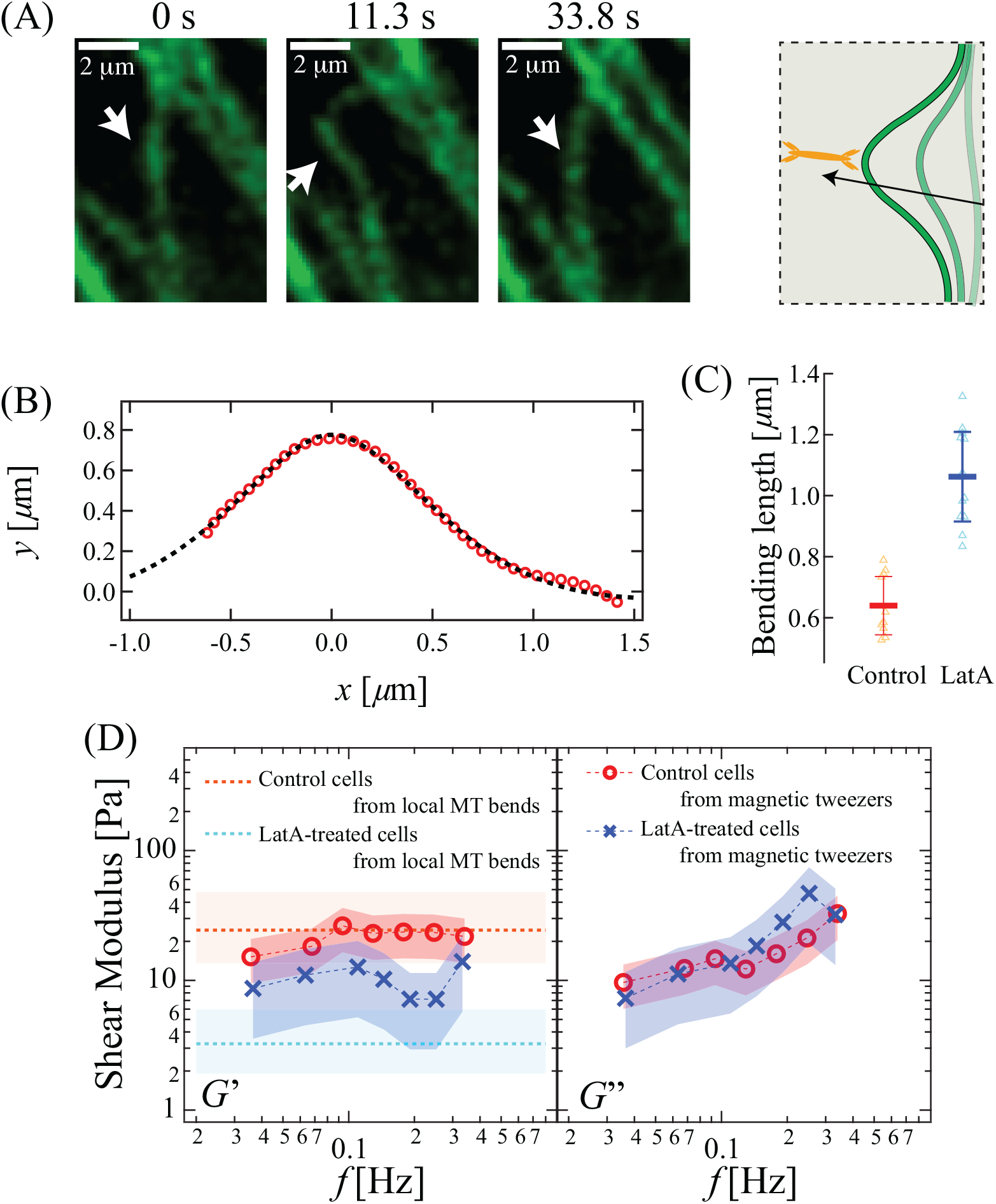
Localized microtubule bends in living cells. **(A)** Snapshots of a microtubule showing a transient localized bend in a Hela cell stained with a SiR-tubulin (see Supplementary movie S3), together with a schematic image. The microtubule was initially straight and relaxed back to a straight shape after a transient deformation. **(B)** Most strongly bent shape of a section of the microtubule shown in (A) (red dots), with a fit of the predicted shape. **(C)** Characteristic axial bending length of localized microtubule bends in HeLa cells with and without Latrunculin A treatment. We tracked 11 microtubules and 13 microtubules for cells without and with Latrunculin A, respectively. Error bars are ± SD. **(D)** Real (*G’*) and imaginary part (*G”*) of the complex shear modulus of the HeLa cell cytoplasm with and without Latrunculin A treatment, measured by magnetic tweezers. Storage moduli estimated from the axial extent of localized microtubule bends are plotted as horizontal dotted lines in the left panel. Error bars are ± SD.

After establishing and testing our new method, we set out to investigate how glutamylation, a posttranslational modification of microtubules that is highly abundant in complex microtubule assemblies such as those found in axons, cilia and flagella affects the mechanical properties of microtubles *in vivo*. Polyglutamylation functionalizes the intrinsically disordered C-terminal tails of tubulin that decorate the outside of the microtubule. Glutamylation does not chemically stabilize microtubules directly [60], but this modification is strongly enriched on cellular microtubules that are resistant to temperature and drug-induced depolymerization. Its effects on microtubule mechanics are unknown [6]. Interestingly, glutamylation is strongly enhanced in microtubule arrays that are subject to high mechanical stress such as those in axons and beating cilia. To investigate the effects of glutamylation on microtubule mechanics, we transfected Hela cells with TTLL6-YFP, a polyglutamylase that adds long glutamate chains to *α*-tubulin [61]. TTLL6 is enriched in cilia and is important for the biogenesis and function of this organelle [62, 63]. HeLa cells had low levels of glutamylation in the absence of the overexpressed enzyme (Fig 5A). We confirmed successful polyglutamylation in cells overexpressing TTLL6-YFP by immunofluorescence with the GT335 antibody that recognizes posttranslationally added glutamates (Fig. 5A).

**Figure 5.**
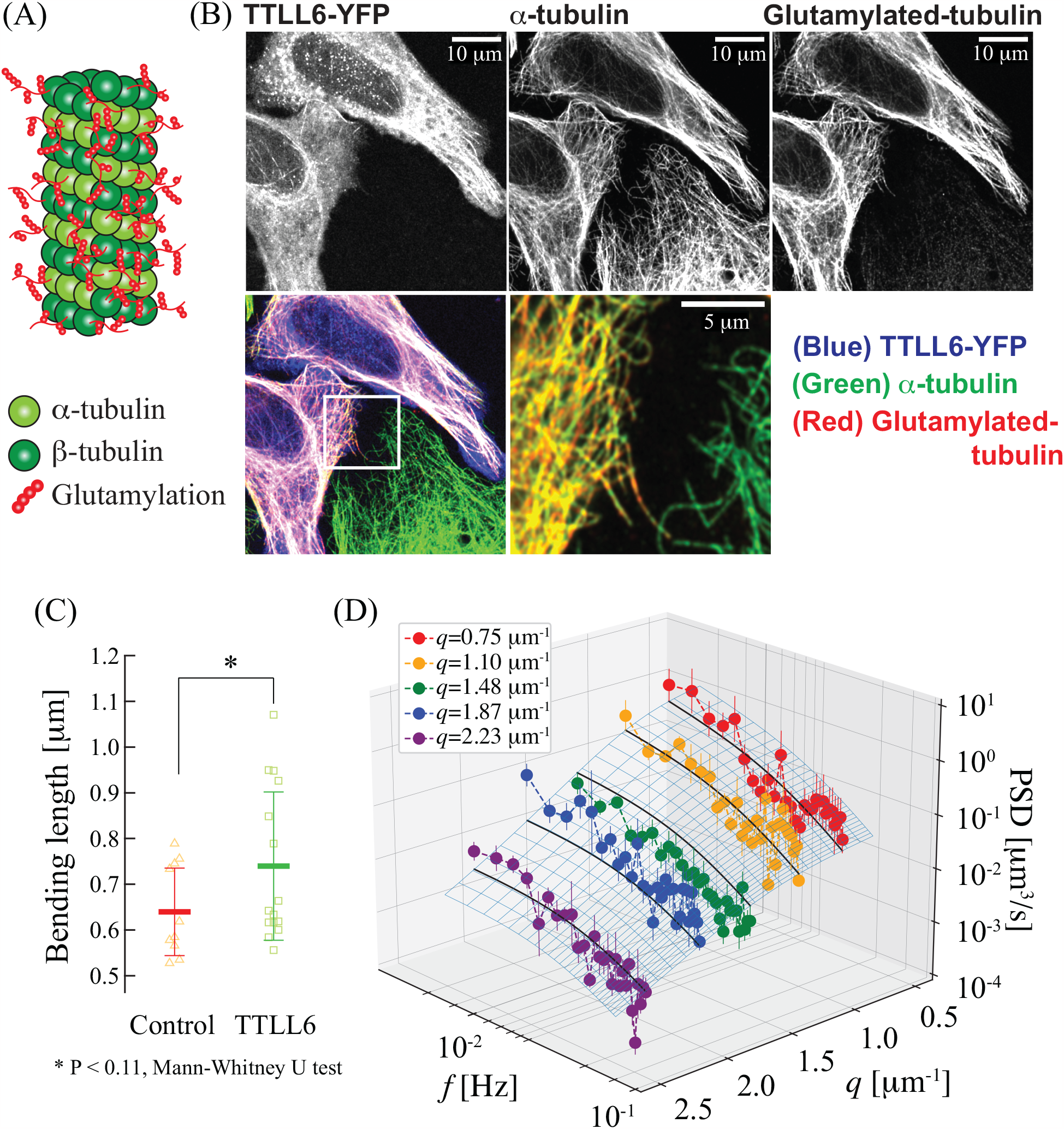
Bending stiffness of polyglutamylated microtubules. **(A)** A schematic illustration of tubulin glutamylation. **(B)** Immunofluorescence images of HeLa cells expressing TTLL6-YFP. Top row: individual fluorescence channels. Bottom row: superposition of images for TTLL6-YFP (blue), microtubules (green) and glutamylated microtubules detected by GT335 (red). **(C)** Bending lengths of localized microtubule bends in HeLa cells expressing TTLL6. (see Supplementary movie S4) The asterisk represents significant difference from the control group based on a Mann-Whitney U test (P < 0.11). **(D)** Global fit of PSDs of mode amplitude fluctuations of microtubules in HeLa cells expressing TTLL6. Results from 15 microtubules were smoothed by binning at *q* = 0.75, 1.10, 1.48, 1.87, and 2.28 *μ*m^-1^. Error bars are ± SEM, bin widths ∼ 0.3 *μ*m^-1^. A 2D fitting plane using Eq. (6) is plotted as a gray mesh plane. Black solid lines represent slices of the fitted plane at *q* = 0.75, 1.10, 1.48, 1.87, and 2.28 *μ*m^-1^, respectively.

To find out if polyglutamylation affects microtubule bending stiffness, we used both methods described above, fluctuation mode analysis and shape analysis of localized bends. We stained microtubules with SiR-tubulin in TTLL6-YFP-transfected cells and imaged strong localized bends. In cells with strong expression of TTLL6-YFP, we observed binding of TTLL6 to microtubules, which may suppress their bending dynamics. We thus selected cells with medium expression of TTLL6-YFP in which no TTLL6 localization was visible on microtubules (Fig. S3). We found that the axial length of localized bends increased to 0.74 (± 0.16) *μ*m in TTLL6-YFP expressing cells, compared with 0.64 (± 0.09) *μ*m in control cells as shown in Fig. 4C. This result is significant (P < 0.11) based on a Mann-Whitney U test. From this result, we estimated the bending stiffness of polyglutamylated microtubules, *κ*_TTLL6_, using the ratio of the cytoplasm elasticities measured in control cells (*G*’_con_) and transfected cells (*G*’_TTLL6_) applying the relation, κ_TTLL6_ ≅ κ_con_*G*_TTLL6_ *l*_TTLL6_ ^4^/(*G*_con_ *l*_con_ ^4^), which is derived from Eq. (8). Our shape analysis shows *l*_TTLL6_ ^4^/*l*_con_^4^ ≈ 1.79. We determined the ratio *G*_TTLL6_/*G*_con_ using active magnetic-bead microrheology (Fig. S4). These results indicate that the bending stiffness of polyglutamylated microtubules (7.8 ×10^-23^ Nm^2^) was significantly larger than that found in control cells for the control microtubules (4.1 ×10^-24^ Nm^2^).

We also tracked bending fluctuations of microtubules in transfected cells and analyzed the mode amplitude PSDs by fitting five scale-dependent PSDs evaluated from 15 microtubules, with a single 2D plane defined by Eq. (6) (Fig. 5C). We again fixed the memory function α(ω) = *k*_0_*G*(ω) in the fit using magnetic-bead rheology results (Fig. S4) on TTLL6-YFP transfected cells. We found *κ* = 7.0 (± 1.7) ×10^-24^ Nm^2^,τ_M_ = 6.2 (± 1.1) s, and *C*_FF_ = 6.4 (± 2.4) ×10^1^ pN^2^/*μ*m to result in a good fit of the data. Analysis of both localized bends and the scale-dependent PSDs thus suggests that polyglutamylated microtubules are ∼ 1.7 times stiffer than microtubules in control cells. Recently, K. P. Wall *et al*. reported that the bending stiffness of *in vitro* reconstituted microtubules increases with polyglutamylation due to the interactions of the *α*-tubulin C-terminal tail (CTT) with the tubulin body, supporting our *in vivo* measurement [11], although it is likely that additional *trans* effects are at play in the cellular situation due to changes in the recruitment of microtubule-binding proteins on polyglutamylated microtubules [7, 64-67].

## Conclusions

We show how quantitative measurements of non-equilibrium shape fluctuations can be used to measure material properties of individual microtubules in living cells. The bending stiffness of microtubules that we measured *in vivo* lies in the window of values reported from *in vitro* studies.Our *in vivo* measurement is non-invasive and requires only that microtubules be fluorescently labeled. A direct characterization of microtubules in living cells opens up possibilities to study microtubules in their native environment and locally track functional changes and variations in their mechanical properties in cells and tissues. After determining microtubule bending stiffness we show how microtubules can be used as an endogenous mechanical probe for cytoplasm mechanics. This method completely avoids the introduction of foreign mechanical probes into cells. We find that the elastic properties of the cytoplasm measured from highly localized bends are in good agreement with those measured by active magnetic-bead mirorheology both for control and latrunculin A-treated cells. Our analysis on both highly localized bends and scale-dependent PSDs of bending amplitudes shows that polyglutamylated microtubules are stiffer than unmodified ones. Such an increased mechanical stability is consistent with the abundance of glutamylated microtubules in stable cellular structures such as the axons of neurons and the axonemes of cilia and flagella. Interestingly, acetylation, a tubulin modification also associated with stable microtubule structures, decreases microtubule stiffness *in vitro* [68], while another study found that microtubule acetylation enhances overall cell stiffness [69]. Our method opens up the possibility to comprehensively study the effects of these modifications on microtubule mechanics in living cells. Furthermore, our method is applicable to other rod-shaped subcellular components embedded in the cytoplasm, for example mitochondria or the endoplasmic reticulum, which are difficult to reconstitute *in vitro* in similar geometries to those they exhibit in cells.

## Acknowledgements

We thank C. P. Broedersz for helpful discussion. A.R-M. is supported by the intramural programs of the National Institute of Neurological Disorders and Stroke (NINDS) and the National, Heart, Lung and Blood Institute (NHLBI). CFS and FCM would like to thank the Isaac Newton Institute for Mathematical Sciences for support and hospitality during the programme New statistical physics in living matter: non equilibrium states under adaptive control. FCM was supported in part by the National Science Foundation Divisions of Materials Research (Grant No. DMR-2224030) and Physics (Grant No. PHY-2019745). RF and KN acknowledge financial support from the Alfred P. Sloan Foundation grant (Grant No. G-2021-14197).

## Author contributions

KN, ARM, and CFS conceived the project. KN and FCM developed the model. KN, RF, ARM and CFS designed the experiments. KN, SR, and AP performed the experiments. KN analyzed and visualized the data. All authors contributed to the writing of the paper.

## Supporting information

## Materials and Methods

### Cell culture

HeLa cells were procured from ATCC, USA (American Type Culture Collection, Manassas, VA, Cat. No. CCL2). Cells were cultured using Dulbecco’s Modified Essential Medium (Thermo Fisher Scientific Inc., Waltham, MA, USA, Cat. No. 11885084) supplemented with 1X Penicillin-Streptomycin (Thermo Fisher Scientific Inc., Waltham, MA, USA, Cat. No. 15140122) and 5% (v/v) Fetal Bovine Serum (Sigma-Aldrich, St. Louis, MO, USA, Cat. No. F2442) inside a sterile and humid 37C incubator with 5% CO2 supply. Cells were passaged using 0.05% trypsin-EDTA (Thermo Fisher Scientific Inc., Waltham, MA, USA, Cat. No. 25300062) when cell density reached ∼90% confluency.

### Staining and imaging microtubules in living cells

HeLa cells were plated on fibronectin (Sigma-Aldrich, St. Louis, MO, USA) coated glass bottom petridishes (ibidi GmbH, Germany, Cat. No. 81158). After ∼3 hours. SiR tubulin (Cytoskeleton, Inc., Denver, CO, Cat. No. CY-SC002) dye (250 nM in DMEM) was applied for live-staining microtubules. After one hour of staining, dye-containing medium was removed, fresh medium was added and cells were imaged under a Zeiss LSM 880-Airy scanning microscope (Zeiss, Germany) using a 63X, NA 1.4 oil objective. The sample chamber was equipped with a CO_2_ supply unit and a temperature controller to achieve 5% CO_2_ concentration and 37 °C temperature inside the chamber. A 633 nm HeNe laser was used to excite the labeled microtubules, and fluorescence emission signals were collected using the appropriate filter in the Airy scanning array detector. Timelapse recordings were taken with intervals of 2-4 s, keeping the z-plane fixed by an IR-based autofocusing unit (Definite Focus).

### Latrunculin A treatment

Culture medium (DMEM) with 1 *μ*M latrunculin A (Sigma-Aldrich, St. Louis, MO, USA, Cat. No. 428021) was used to depolymerize the actin network inside live HeLa cells. Timelapse recordings of SiR-tubulin labeled microtubules were acquired until up to 1 hr after addition of Latrunculin A (LatA). The time window for observation was optimized by checking the effect of Lat-A on actin. After 1 hr of treatment, cells started to round up (see supplementary information). Therefore, we imaged microtubule only up to 1 hr after treatment with LatA.

### Transfecting cells with TTLL6-YFP and imaging

TTLL6-YFP [70] was transiently expressed in HeLa cells using a Lipofectamine transfection kit (Thermo Fisher Scientific Inc., Waltham, MA, USA, L3000008). Cells were then incubated with SiR-tubulin to label microtubules after 8-10 hours post transfection. TTLL6-YFP expression and microtubule dynamics were imaged using the same microscope (Zeiss 880) with same settings (as described above in the section under ‘Staining and imaging microtubules in living cells’) for timelapse imaging. We used the 514 nm laser line of an Argon gas laser to excite YFP.

### Image Analysis

The coordinates of the backbones of microtubules were determined from each image by the JFilament plugin in ImageJ [71] and used for further data processing. After extracting the backbone coordinates (*x*_*i*_,*y*_*i*_) of microtubules, the amplitude *a*_*q*_(*t*) of the *n*th mode was calculated from the local tangent angle of the filament [12] *θ*(*s*_*i*_) = tan^-1^(*yi*_+1_-*y*_*i*_)/(*xi*_+1_-*x*_*i*_) via integration by parts 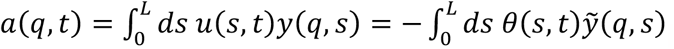, based on the relation 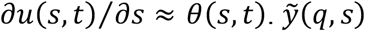 denotes the integral of *y*(q, *s*). The orthogonal eigenmodes are given by *y*(*q*,*s*) as 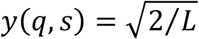 sin q*s*, as discussed in the main text.

### Antibody staining and imaging of fixed cells

HeLa cells were fixed using 4% paraformaldehyde and permeabilized using Triton-X100 (VWR, Radnor, PA, USA, Cat. No. M143-1L) 12 hours after TTLL6-YFP transfection. To confirm that tubulin glutamylation occurred in the TTLL6-transfected cells, we used a primary antibody specific for glutamylated microtubules (1:5000, AdipoGen Life Sciences, Inc. Cat. No. mAB GT335) and a Cy5-tagged secondary antibody (1:2000, Thermo Fisher Scientific Inc., Waltham, MA, Cat. No. A10524). Total tubulin was labeled using a primary antibody against alpha-tubulin and a Cy3-tagged secondary antibody (Thermo Fisher Scientific Inc., Waltham, MA, USA).

After staining cells, PBS was added and fixed cells were imaged under the Zeiss 880 microscope using Airy Scanning mode and a 63 X, NA 1.4 oil immersion objective. The 514 nm laser line from an Argon gas laser, a 561 nm Diode Pumped Solid State Laser and a 633 nm HeNe gas laser were used for imaging TTL6-YFP, alpha-tubulin and glutamylated tubulin, respectively.

### Magnetic tweezers and force calibration

The magnetic tweezers setup consists of a brass solenoid fitted on a cylindrical core made of low-carbon steel AISI 1006. The low-carbon steel was selected to maximize magnetic flux generated by the core due to its high magnetic induction compared to other commonly used alloys in magnetic tweezers such as MuMetal (Ni-Fe Alloys) [72, 73]. The solenoid has a length of 50 mm and an outer diameter of 20 mm, and is wound on a brass frame with 1000 turns of 24 AWG copper wire. The magnetic core has a diameter of 5 mm and a length of 150 mm. One end of the core is tapered to a tip with an angle of 60 degrees and a tip radius of <10 *μ*m. The sharp tip ensures a high magnetic field gradient, maximizing the force on paramagnetic beads. The magnet is held by a Leica mechanical micromanipulator (Leica, Germany) and is manually adjusted to the minimal possible distance to the cell (∼50 μm). A customized Matlab code is used to generate voltage signals via a data I/O card (NI USB-6001, National Instruments). Those signals drive a high-current amplifier (Kepco Inc., Flushing, NY, USA, Model: BOP 36-6M) the output of which is connected to the solenoid. The magnetic tweezers setup was mounted on an inverted confocal microscope (SP5, Leica, Germany) equipped with a 63x 1.4 NA objective HCX PL APO, Leica, Germany). Bright field images of beads were recorded with a CCD camera (UI-3240CP-M-GL, IDS, Germany) with the frame rate of 4 Hz.

To calibrate the force exerted by the magnetic tweezers, 1 *μ*m paramagnetic beads were dispersed in a viscous liquid of known viscosity. Here, we used poly-dimethylsiloxane (PDMS) oil (Sigma-Aldrich, St. Louis, MO) with a dynamic viscosity of 9.71 Pa·s. We positioned the microscope stage and the magnetic tweezers such that a single bead was positioned 50 *μ*m in front of the needle tip. Then we applied a constant current to the solenoid and tracked bead displacement. From the bead velocity we calculated the force acting on the particle according to Stokes’ law: *F* = 6*prhv*, where *r* is the radius of the bead, *h* is the viscosity of the oil, and *v* is the bead velocity. The measured force vs. distance relationship (Fig. S2) followed a stretched exponential function with respect to the bead-needle distance. We used these fitting results for further analysis.

### Preparation of PEG-grafted beads

Amine-terminated methoxy-poly(ethylene glycol), NH2-(CH2-CH2-O)n-OCH3 (mPEG-NH2) with average n = 17 and average molecular mass 750 Da, 2-(N-morpholino) ethanesulfonic acid (MES), and 1-[3-(dimethylamino)propyl]-3-ethylcarbodiimide (EDC) were purchased from Sigma Aldrich Corp. (St. Louis, MO, USA). Carboxylated 1 *μ*m paramagnetic beads were obtained from Thermo Fischer (Dynabeads MyOne Carboxylic Acid, ThermoFisher, USA). 5 *μ*l of the bead solution were centrifuged, and the supernatants were discarded. Beads were resuspended in 500 *μ*l of 0.1 M pH5 MES buffer (VWR, Radnor, PA, USA, Cat. No. E149-100G). 12.5 *μ*l of the 0.2 g/ml EDC solution and 50 *μ*l of the 1 mg/ml mPEG-NH_2_ solution were then added to the bead solution and incubated overnight at 4 °C. After the reaction, the bead solution was centrifuged and the supernatant was discarded. Beads were resuspended in water and stored as a stock solution for up to three months at 4 °C.

### Sample preparation for active magnetic-bead microrheology

Transfection for GFP and insertion of magnetic beads was performed using a 4D-Nucleofector (Amaxa Biosystems) by optimizing a protocol for HeLa cells (solution SE, program CN114). In each nucleofection experiment, the paramagnetic bead solution and pmaxGFP vector were added. After the reaction, the nucleofected cells were immediately transferred into fresh medium in Petri dishes (μ-Dish 35 mm, ibidi, Germany, Cat. No. 80136) and incubated for 24 - 48 h.

### Cellular ATP depletion

2-deoxy-glucose (2-DOG) (Sigma-Aldrich, St. Louis, MO, USA, Cat. No. D3179) was used to block ATP production through the glycolytic pathway and Oligomycin-A (OA) (Sigma-Aldrich, St. Louis, MO, USA, Cat. No. 75351) was used to block ATP production by mitochondria. After staining the microtubules by SiR-tubulin (250 nM), HeLa cells were treated with glucose-free DMEM (supplemented with 10X FBS and 1X PenStrep) containing 25 mM of 2-DOG and 2.5 *μ*M OA. Intracellular movement started to slow down after the treatment. After one hour, we did not observe any more significant fluctuations of microtubules.

### Estimation of errors in the fitting of the scale-dependent PSDs

There are two major sources of experimental uncertainties: (i) Random variations in the PSDs measured from different microtubules due to finite recording times, microscope resolution and finite tracking accuracy and differences in the microtubules and their local environments, (ii) variations in the complex shear modulus measured by active magnetic-bead MR due to finite image resolution and tracking accuracy and variations in the local bead environments. We used a bootstrapping method to estimate the resulting uncertainties in the fitting parameters of the scale-dependent PSDs. We computer-generated 100 additional PSDs and 100 magnetic bead response curves by adding random numbers to the average experimental PSDs and magnetic bead response curves, generated from normal distributions with a mean of 0 and a standard deviation equal to the observed experimental s.e.m.

We then fitted the PSDs using Eq. 6. with three fitting parameters, namely *C*_FF_, τ_M_, and κ. In order to maximize the accuracy of the error estimates, we first used asymptotic form of Eq. 6, valid in the limit of *q* ≪ (α(ω)/κ)^1/4^ and *ω* = 2*πf* ≪ 1/τ_M_:

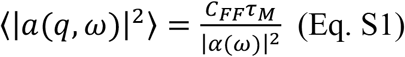

In this limit, the microtubule shape fluctuations are only marginally influenced the bending stiffness and are simply determined by the balance between active forces from molecular motors (*C*_FF_) and the cytoplasm viscoelasticity (α(ω)). Using the literature values, *G*(ω)∼ 10 Pa, κ ≲ 1×10^-23^ Nm^2^, and *π*_M_ ∼ 10 s [12, 25, 36, 37], we applied Eq. S1 to PSDs within the range of *q* < 1.1 *μ*m^-1^ and *f* < 0.015 Hz to determine the term *C*_FF_*π*_M_. We subsequently fitted Eq. 6 to all complete data sets to determine the other two parameters. Histograms of fitting parameters are summarized in Fig. S5.

Movie S1: Non-equilibrium bending fluctuations of microtubules stained with SiR-tublin in a Hela cell, acquired at 0.4 fps (sped up 10x) (scale bar: 1 *μ*m)

Movie S2: A localized microtubule bend in a Hela cell stained with SiR-tublin, acquired at 0.4 fps (sped up 5x) (scale bar: 1 *μ*m)

Movie S3: Non-equilibrium bending fluctuations of microtubules in a Hela cell expressing TTLL6-YFP stained with SiR-tublin, acquired at 0.4 fps (sped up 5x) (scale bar: 3 *μ*m)

Movie S4: A localized microtubule bend in a Hela cell expressing TTLL6-YFP stained with SiR-tublin, acquired at 0.4 fps (sped up 5x) (scale bar: 2 *μ*m)

**Figure S1.**
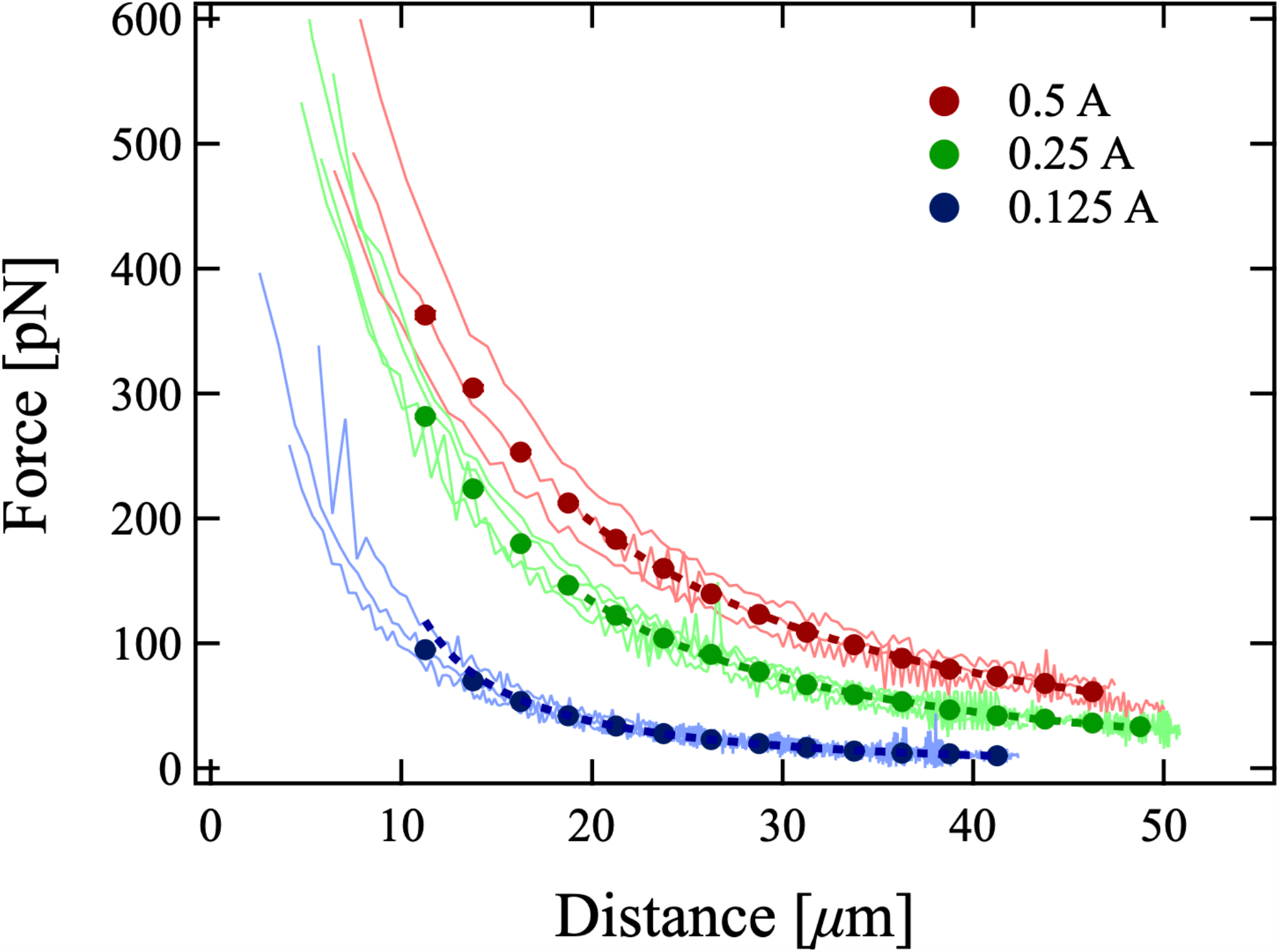
Calibration of magnetic tweezers. Calibration is performed by tracking bead velocity in high-viscosity PDMS oil (Sigma-Aldrich, St. Louis, MO) with a dynamic viscosity of 9.71 Pa·s, as a function of distance to the tip. Force-distance curves for different currents. Solid lines indicate the fits with a stretched exponentials.

**Figure S2.**
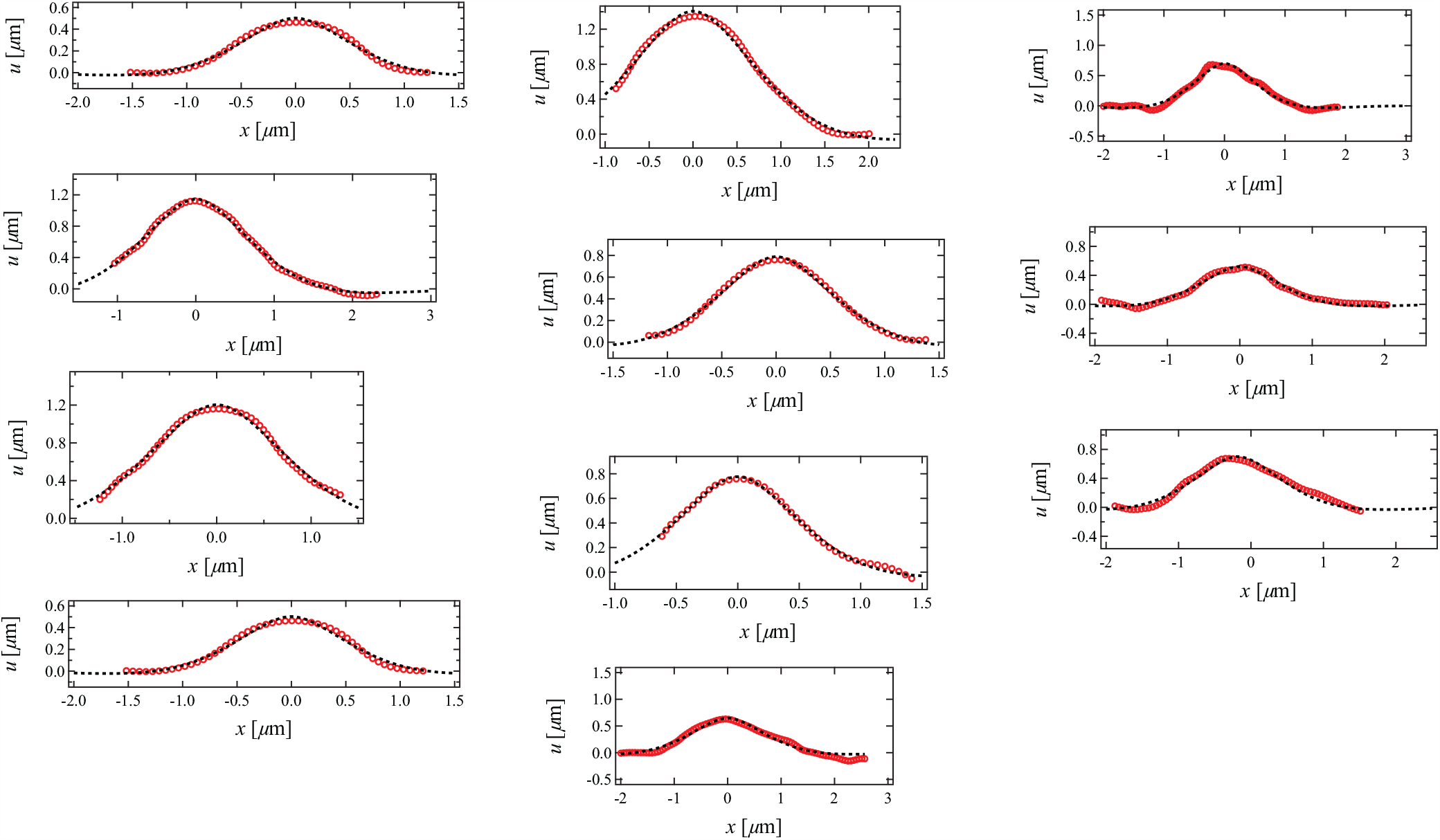
Fitting the shapes of strongly bent microtubules. Tracked shapes of highly localized bends in living HeLa cells (red lines) and fits with the predicted form (Eq. 9) (black dots). We found good agreement of the observed microtubule shapes with Eq. 9. From these fits, we extracted the bend length *l* as summarized in Fig. 4C.

**Figure S3.**
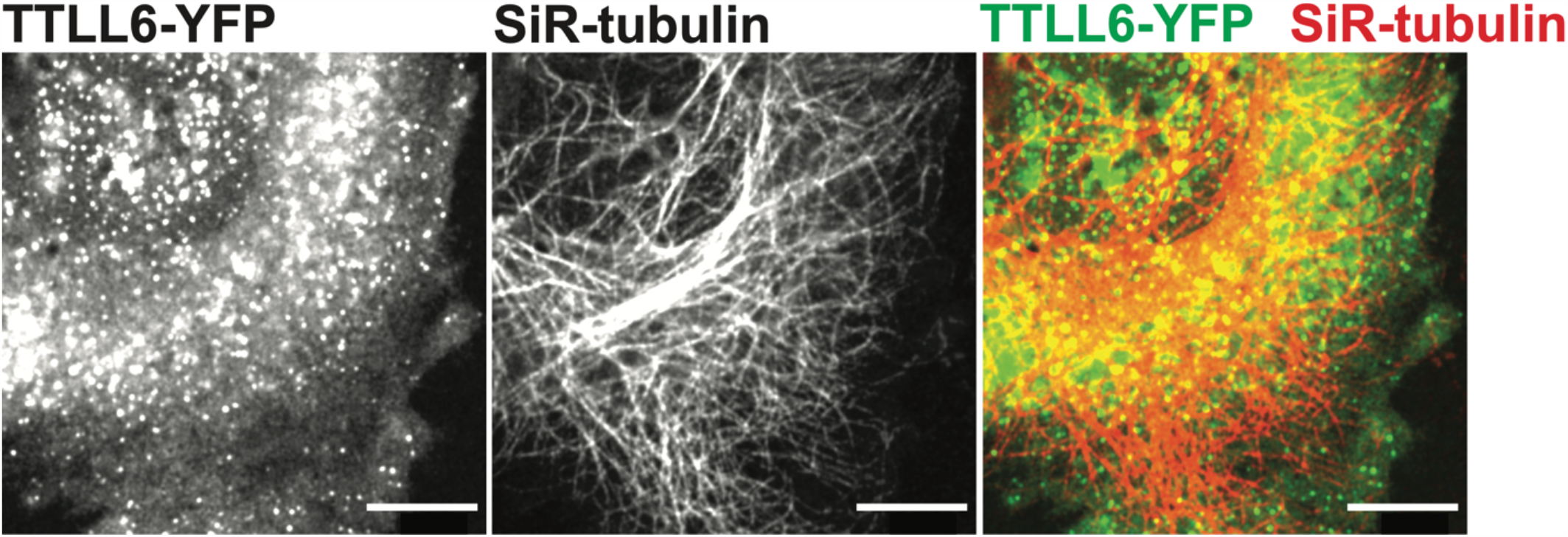
Expression of TTLL6-YFP. Representative images of living HeLa cells expressing TTLL6-YFP, stained for microtubules with SiR-tubulin, individual channels and superposition (scale bar: 10 *μ*m).

**Figure S4.**
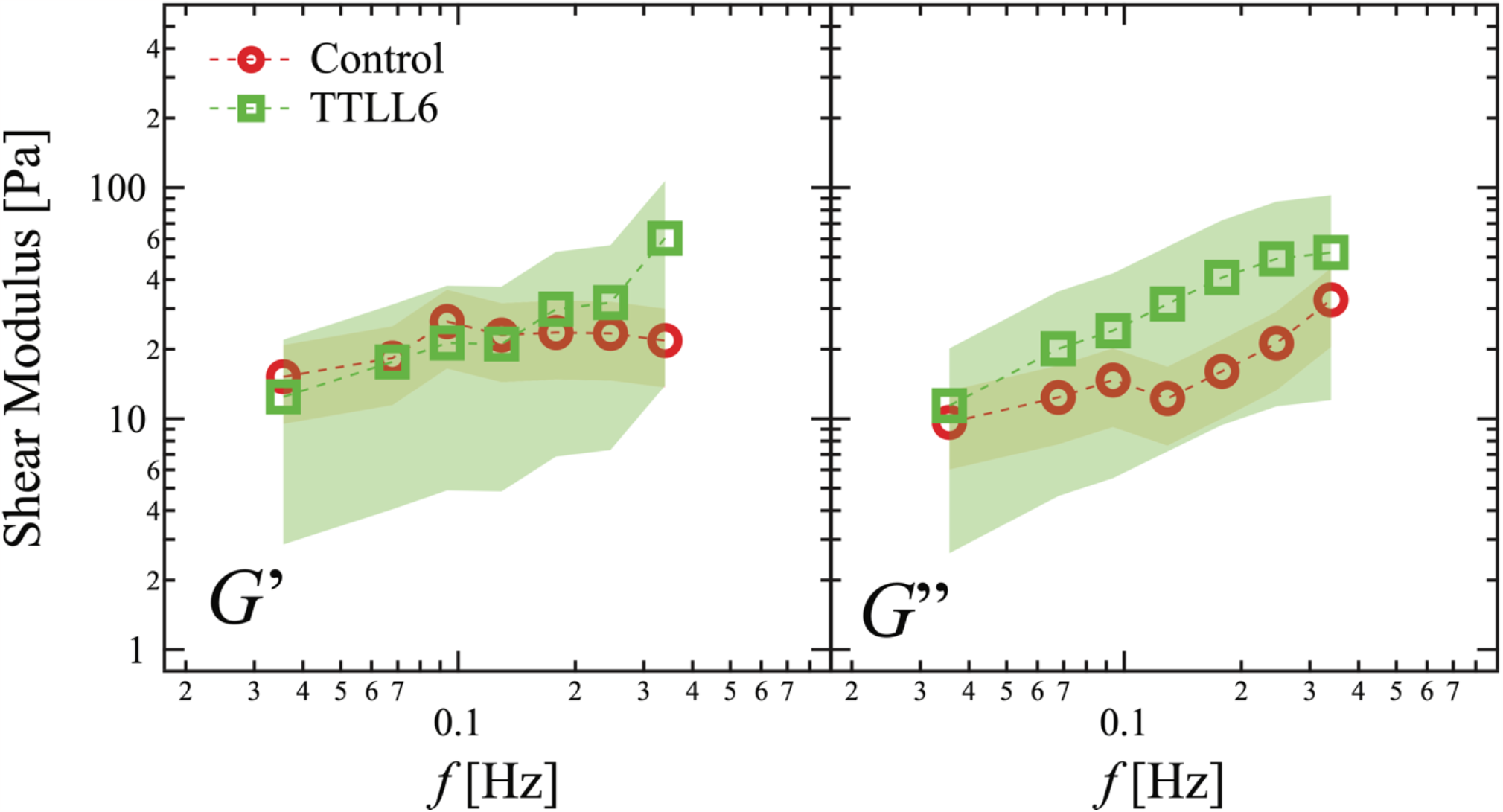
Magnetic tweezers microrheology in TTLL6-YFP-expressing cells. Frequency-dependent shear moduli were evaluated from the creep compliance *J*(*t*) through Eq. (7) for control cells (*N* = 9) and TTLL6 expressing cells (*N* = 14), with real part or storage modulus *G*’ and imaginary part or loss modulus *G*”. Error bars reflect ± SD. As discussed in Fig. 5, we evaluated the ratio of the cytoplasm elasticities measured from control cells (*G*’_con_) and TTLL6 expressing cells (*G*’_TTLL6_) to estimate the bending stiffness of polyglutamylated microtubules, _TTLL6_. *G*’ of both control cells and TTLL6 expressing cells were fitted by a constant function around *f* = 0.1 - 0.2 Hz to determine *G*_con_ and *G*_TTLL6_.

**Figure S5.**
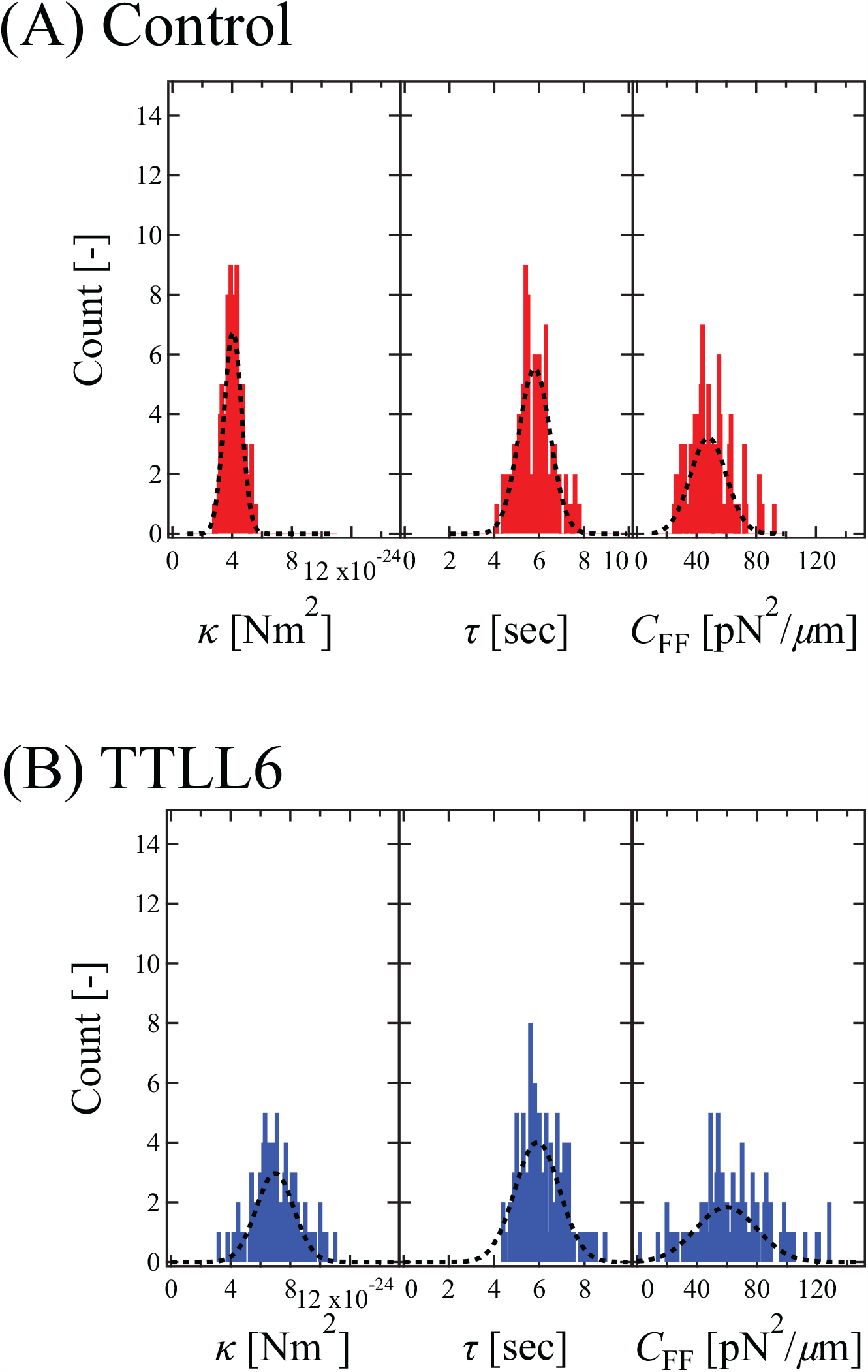
Histogram of fitting parameters obtained by a global fit of PSDs with Eq. (6) on in (A) control HeLa cells and (B) cells expressing TTLL6-YFP. Black dotted lines are Gaussian fitting results.

